# Expression of NMNAT1 in the Photoreceptors is Sufficient to Prevent *NMNAT1*-Associated Disease

**DOI:** 10.1101/2022.11.20.517250

**Authors:** Emily E. Brown, Michael J. Scandura, Eric A. Pierce

**Author notes:** Correspondence should be addressed to E.A.P.

## Abstract

Nicotinamide nucleotide adenylyltransferase 1 (NMNAT1) is a ubiquitously expressed enzyme involved in nuclear NAD^+^ production throughout the body. However, mutations in the *NMNAT1* gene lead to retina-specific disease with few reports of systemic effects. We have previously demonstrated that AAV-mediated gene therapy using self-complimentary AAV (scAAV) to ubiquitously express NMNAT1 throughout the retina prevents retinal degeneration in a mouse model of *NMNAT1*-associated disease. We aimed to develop a better understanding of the cell types in the retina that contribute to disease pathogenesis in *NMNAT1*-associated disease, and to identify the cell types that require NMNAT1 expression for therapeutic benefit. To achieve this goal, we treated *Nmnat1^V9M/V9M^* mice with scAAV using cell type-specific promoters to restrict NMNAT1 expression to distinct retinal cell types. We hypothesized that photoreceptors are uniquely vulnerable to NAD^+^ depletion due to mutations in *NMNAT1*. Consistent with this hypothesis, we identified that treatments that drove NMNAT1 expression in the photoreceptors led to preservation of retinal morphology. These findings suggest that gene therapies for *NMNAT1*-associated disease should aim to express NMNAT1 in the photoreceptor cells.

## Introduction

Mutations in the *NMNAT1* gene lead to severe early onset inherited retinal degeneration (1). Although this gene is expressed ubiquitously, mutations in *NMNAT1* primarily lead to retina-specific disease, with few reports of systemic disease (1–5). In a small number of patients, structural variant (SVs) in *NMNAT1* have been reported to lead to systemic effects manifest with a phenotype known as known as SHILCA, which is characterized by spondylo-epiphyseal dysplasia, sensorineural hearing loss, and intellectual disability, in addition to the LCA phenotype (2, 6). Interestingly, patients with mutations in *NMNAT1* have atrophy of the macula (1), the region of the retina with a high density of cone photoreceptors, which are responsible for high acuity central vision. There are currently no FDA-approved therapies for *NMNAT1*-associated disease, and the mechanisms underlying the retina-specific nature of this disease remain poorly understood.

Nicotinamide nucleotide adenylyltransferases (NMNATs) are enzymes involved in NAD^+^ biosynthesis. There are three different *NMNAT* genes that encode different NMNAT enzymes that localize to different subcellular compartments. NMNAT1 is localized to the nucleus and is responsible for nuclear production of NAD^+^, NMNAT2 is localized to the cytoplasm and Golgi, while NMNAT3 is localized to the mitochondria. Although it is well characterized that mutations in *NMNAT1* lead to retina-specific disease, the association of variants in other NMNAT isoforms with disease is less understood. Variants in *NMNAT2* have been associated with polyneuropathy with erythromelalgia (7), while GWAS studies have suggested that variants in *NMNAT3* are associated with Alzheimer’s disease (8). More work is needed to better understand the genetic causality of variants in the *NMNAT2* and 3 genes.

We have previously characterized a mouse model of *NMNAT1*-associated disease with the hypomorphic p.V9M mutation in *Nmnat1* (*Nmnat1^V9M/V9M^*) (9). Although the retinas of these mice develop normally, they exhibit progressive retinal degeneration, with reduced retinal function and loss of photoreceptor cells by 4 weeks of age (9). Using human recombinant protein, we found that the p.V9M mutation in *NMNAT1* results in a 63% reduction in NMNAT1 enzymatic activity, suggesting that reduced NMNAT1 function is sufficient to drive disease pathology (1). Knockout of *NMNAT1* is embryonic lethal, indicating that at least some enzymatic function is required for normal development (10).

We investigated the molecular mechanisms of disease pathogenesis of *NMNAT1*-associated disease using the *Nmnat1^V9M/V9M^* model, and have shown that NAD^+^ levels are reduced in the neural retina of these mice, but not in other tissues, prior to retinal degeneration (11). We found that the retinas of *Nmnat1^V9M/V9M^* mice exhibit photoreceptor-specific DNA damage, which is associated with elevated Poly (ADP-ribose) polymerase (PARP) enzymatic activity, suggesting a unique requirement of nuclear NAD^+^ in photoreceptor cells (12). PARP1 is the primary consumer of NAD^+^ in the nucleus and facilitates DNA repair and transcriptional regulation (13), therefore, we hypothesize that photoreceptor cells are uniquely vulnerable to genotoxic stress.

We have previously shown that AAV-mediated delivery of a normal copy of codon optimized human NMNAT1 (hNMNAT1) can prevent retinal degeneration in the *Nmnat1^V9M/V9M^* mouse model of *NMNAT1*-associated disease (14). Using the ubiquitous CASI promoter, we showed that expression of hNMNAT1 in most retinal cell types was sufficient to prevent retinal degeneration and preserve retinal function out to 9 months of age, the latest age examined (14). In the present study, we aimed to determine which cell types in the retina require NMNAT1 expression, to better understand which cell types need to be targeted for successful AAV-mediated gene therapy and which cell types contribute to disease pathology. To this end, we delivered a variety of different constructs with cell type-specific promoters to drive expression of NMNAT1. These included canonical promoters as well as synthetic promoters that have been characterized previously (15). We show that expression of NMNAT1 solely in the RPE is not sufficient to prevent retinal degeneration. However, expression of NMNAT1 in the neural retina is sufficient to prevent retinal degeneration, with the greatest benefit observed with NMNAT1 expression throughout the retina, followed by expression of NMNAT1 specifically in the photoreceptor cells. These findings suggest that gene therapies for NMNAT1-associated disease may be most effective if they aim to drive expression of NMNAT1 in the photoreceptor cells, with potential additional benefit with expression throughout the neural retina.

## Results

Using a mouse model of NMNAT1-associated disease (*Nmnat1^V9M/V9M^*), we have previously shown that AAV-mediated gene augmentation therapy is able to prevent the loss of both retinal structure and function. We identified that the more rapidly expressing self-complimentary AAV (scAAV), rather than single-stranded AAV (ssAAV), is required for protection due to the narrow therapeutic window in this model (14). Early injections prior to postnatal day 12 (P12) do not lead to transduction of the necessary cells in the retina, as the retina is still in development. However, there is early rapid onset retinal degeneration in the *Nmnat1^V9M/V9M^* mice, with loss of photoreceptor cells by 4 weeks of age (P28). Therefore, treatment within a therapeutic window between P12 and P18 is required. We showed that 2 weeks post injection, treatment with a ssAAV results in a lower level of expression of hNMNAT1 than treatment with scAAV (14). The reduced levels of expression with ssAAV are likely due to more rapid expression of the transgene with scAAV, which does not require second strand synthesis (16).

We have previously shown that photoreceptors appear particularly susceptible to NMNAT1-depletion due to mutations in *NMNAT1*, as they display elevated levels of genotoxic stress. Therefore, we hypothesized that photoreceptors are uniquely sensitive to NMNAT1 enzymatic activity, and that photoreceptor-specific expression is sufficient to prevent retinal degeneration. To this end, we delivered a variety of AAV constructs with cell type-specific promoters to the retinas of *Nmnat1^V9M/V9M^* mice. All constructs were delivered using scAAV2/9 with cell type-specific promoters driving the expression of codon optimized hNMNAT1. With the goal of testing promoters with translatability to a human therapy, we selected human-specific promoters or promoters that have been shown to express in the desired cell types in human cells. We also selected promoters based on their length and ability to fit into a scAAV.

All vectors were delivered at 1.0 E9 vector genomes per microliter (vg/μl) and co-administered with scAAV2/9.CMV.EGFP delivered at 1.0E 8 vg/μl. The quality of the subretinal injections was assessed 6 weeks post injection, utilizing a non-invasive procedure called fundus photography that uses a low powered microscope attached to a camera to take images of the back of the eye. Successful injections were confirmed by visualization of the injection site, and the expression of GFP observed using a GFP filter with the funduscope (Figure S1A). Optical coherence tomography (OCT), a non-invasive method that allows visualization and quantification of the thickness of the retinal layers was utilized to assess preservation of retinal structure. We performed OCT measurements in the inferior region of the retina, which is adjacent to the injection site and is likely to be treated with more virus, the superior region of the retina, which is more distal to the injection site, and the inferior region of the untreated contralateral eye of the same mouse (Figure S1B). Both inferior and superior measurements were taken to assess the effect of the treatment beyond the site of the injection, and to determine if there were differences in thickness both near and distal to the injection site. Measurements were taken from the nerve fiber layer (NFL) to the RPE to measure total retinal thickness, and from the outer plexiform layer (OPL) to the RPE to measure thickness of primarily the photoreceptor cell layer (Figure S1B).

We first confirmed our previous finding that pan-retinal expression in the *Nmnat1^V9M/V9M^* model is sufficient to prevent retinal degeneration. Delivery of scAAV2/9.CASI.hNMNAT1.WPRE resulted in expression of NMNAT1 throughout the retina at both 2 and 4 weeks post injection (Figure 1A), as visualized by immunohistochemistry (IHC) of frozen retinal sections stained with an antibody specific to human NMNAT1 (14). We did not observe any positive signal for hNMNAT1 in the untreated contralateral eye for any of the vectors tested (Figure S2). We found that total retinal thickness was significantly preserved in both the inferior and superior regions of the retinas treated with scAAV2/9.CASI.hNMNAT1.WPRE, as compared to the contralateral untreated eye in *Nmnat1^V9M/V9M^* mice (Figure S3). Additionally, the thickness of the photoreceptor nuclear layer, measured as the OPL to the RPE, was significantly greater in the treated eye, as compared to the untreated eye (Figure 2A). These findings confirm our previous results that expression of hNMNAT1 throughout the retinas of *Nmnat1^V9M/V9M^* mice is sufficient to prevent retinal degeneration due to loss of NMNAT1 function.

**Figure 1.**
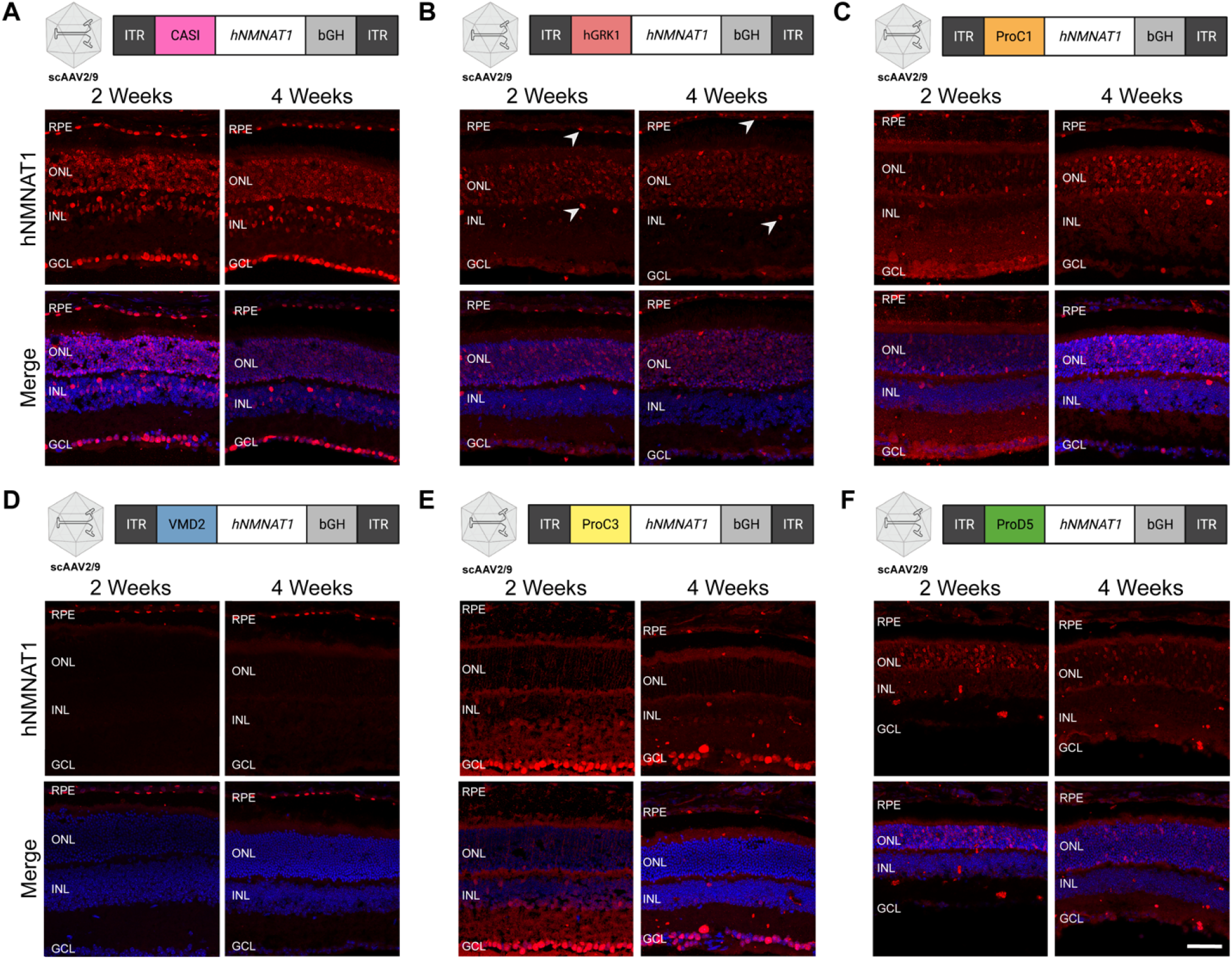
Cell type-specific expression of NMNAT1 using various promoters. Diagrams show the design of AAV constructs with various promoters. Immunofluorescence images demonstrate the cell type-specific expression of hNMNAT1 (red) after delivery scAAV9 at a titer of 1E9 vg/μl, 2- and 4-weeks post subretinal injection. Expression of NMNAT1 using (A) the CASI promoter, which led to expression in most cell types, (B) hGRK1 which led to expression in the photoreceptor cells as well as some expression in RPE and the INL (arrows), (C) ProC1, which led to expression specifically in the photoreceptor cells, (D) VMD2, which led to expression in the RPE, (E) ProC3, which led to expression in the retinal ganglion cells, and (F) ProD5, which led to expression in the rods. Abbreviations: retinal pigment epithelium (RPE); outer nuclear layer (ONL); inner nuclear layer (INL); ganglion cell layer (GCL). Top panels show staining with an antibody specific to human NMNAT1 (hNMNAT1). Bottom panels are counterstained with DAPI (blue). Scale bar indicates 20 μm.

**Figure 2.**
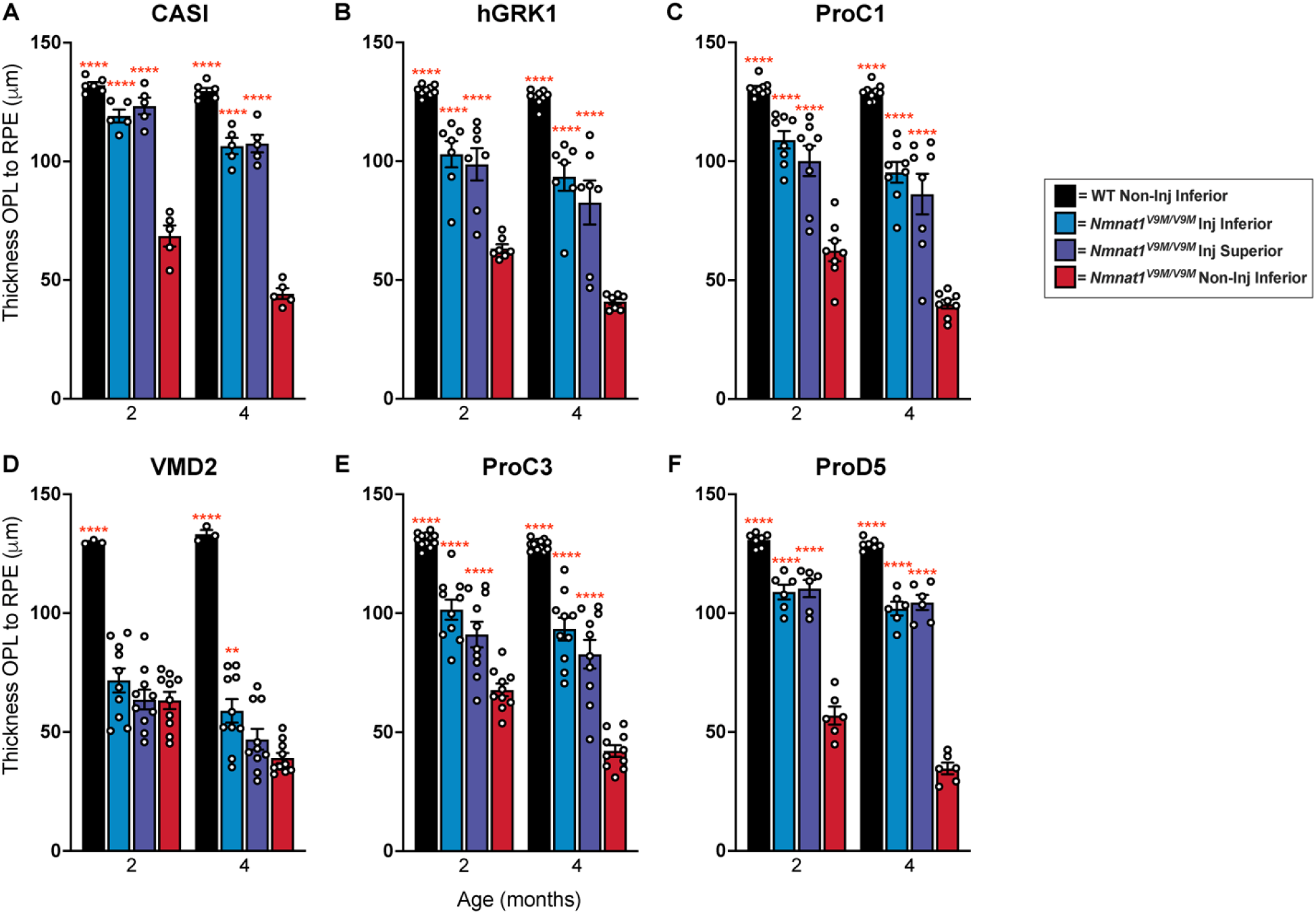
Photoreceptor layer thickness is preserved with expression of NMNAT1 in the neural retina. Measurements of retinal thickness measured from the outer plexiform layer (OPL) to the retinal pigment epithelium (RPE). Measurements are based on SD-OCT imaging of retinas from mice treated with scAAV9 with promoters driving expression of hNMNAT1 in various cell types. The promoters used include (A) CASI (most cell types), (B) hGRK1 (rods and some RPE), (C) ProC1 (photoreceptors), (D) VMD2 (RPE), (E) ProC3 (ganglion cells), (F) ProD5 (rods). Abbreviations: outer plexiform layer (OPL); retinal pigment epithelium (RPE); inner nuclear layer (INL); ganglion cell layer (GCL); spectral-domain optical coherence tomography (SD-OCT). Statistically significant preservation of retinal thickness between treated and untreated *Nmnat1^V9M/V9M^* retinas are noted by red asterisks (n ≥ 3 WT and n ≥ 5 *Nmnat1^V9M/V9M^* mice for each promoter at each age, 2Way ANOVA, multiple comparisons, *P* < 0.05, error bars represent the SEM, points represent the respective retinal thickness for each eye from a single mouse).

Since we have shown that expression of NMNAT1 throughout the retina is sufficient to prevent retinal degeneration in *Nmnat1^V9M/V9M^* mice, and our previous work suggests that genotoxic stress in the photoreceptor cells occurs prior to retinal degeneration, we aimed to assess whether expression only in the photoreceptor cells is sufficient to prevent retinal degeneration in *Nmnat1^V9M/V9M^* mice. To drive expression of NMNAT1 in the photoreceptor cells specifically, we used the human rhodopsin kinase promoter (hGRK1) (17). Treatment with scAAV2/9.hGRK1.hNMNAT1.WPRE resulted in expression in the photoreceptors 2- and 4-weeks post injection, but some expression was also observed in the RPE and the INL (Figure 1B). Expression in the photoreceptors and RPE led to significant preservation of total retinal thickness (P < 0.0001) (Figure S3B) and photoreceptor layer thickness (P < 0.0001) (Figure 2B) in *Nmnat1^V9M/V9M^* retinas, as compared to untreated controls. These findings suggest the expression in the rods, cones, and the RPE is sufficient to significantly prevent retinal degeneration in *Nmnat1^V9M/V9M^* retinas. Expression of NMNAT1 in most cell types in the retina using the CASI promoter appeared to provide a greater benefit (Figure S3, Figure 3, Table 1), although the difference in retinal thickness in the inferior regions of *Nmnat1^V9M/V9M^* retinas was not statistically different between the CASI and hGRK1 promoters. The average NFL to RPE thickness for CASI and hGRK1 were 119.1 μm and 102.93 μm, respectively at 2 months of age, and at 4 months of age were 106.45 μm and 93.5 μm, respectively.

**Figure 3.**
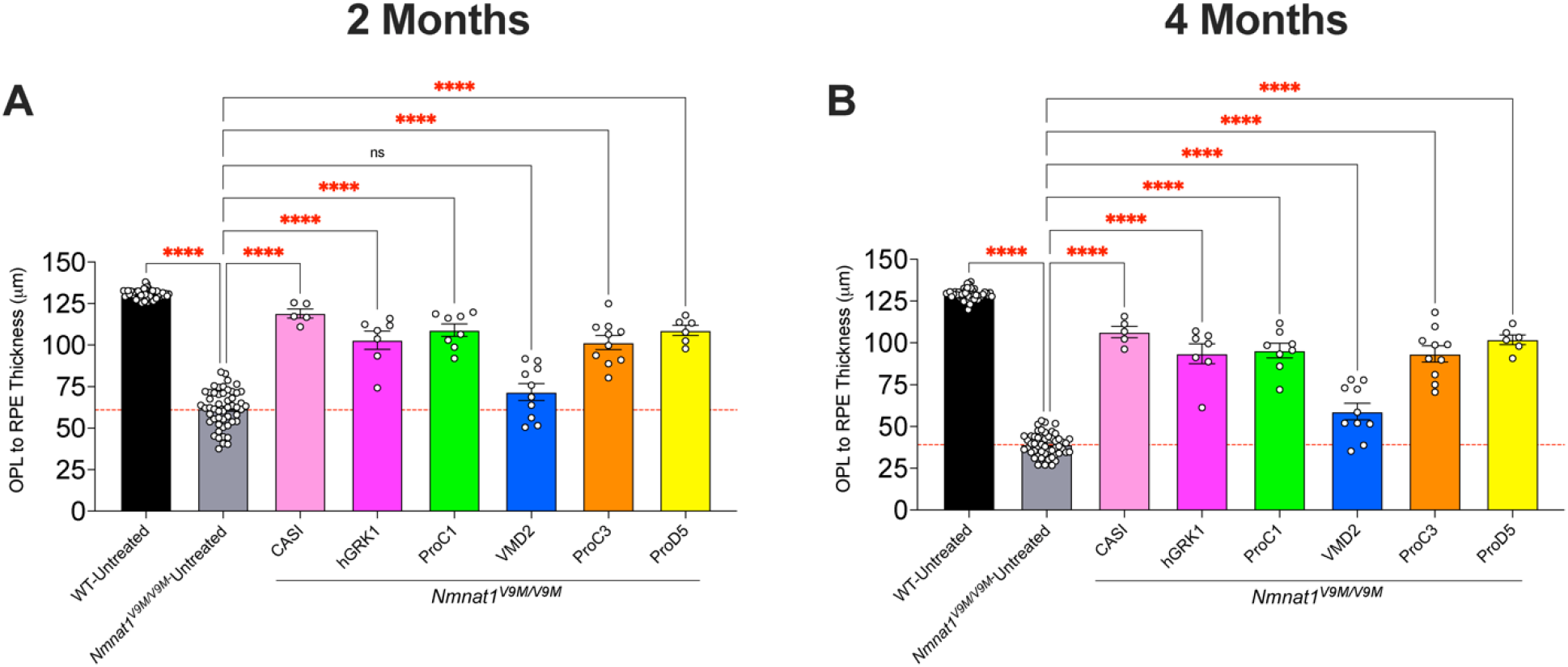
Comparison of preservation of retinal thickness with various promoters. Measurements of retinal thickness measured from outer plexiform layer (OPL) to the retinal pigment epithelium (RPE) at 2 months (A) and 4 months (B) of age. Measurements for *Nmnat1^V9M/V9M^* untreated retinas represent the average retinal thickness for the untreated contralateral eye for all groups. Wild type (WT) untreated measurements represent the average retinal thickness for the untreated contralateral eye from all WT groups. Measurements represent the thickness in the inferior region of the retina (closest to the injection site). All treated eyes were treated with scAAV9 with the respective promoter driving expression of hNMNAT1. This virus was delivered at 1E9 vg/μl and co-injected with AAV9 CMV.GFP at a titer of 1E8 vg/μl. The red dashed line represents the mean thickness of the untreated *Nmnat1^V9M/V9M^* retinas at the respective age. Statistically significant preservation of retinal thickness between untreated and treated *Nmnat1^V9M/V9M^* retinas are noted by red asterisks (n ≥ 3 WT and n ≥ 5 *Nmnat1^V9M/V9M^* mice for each group at each age, 2Way ANOVA, multiple comparisons, *P* < 0.05, error bars represent the SEM, points represent the respective retinal thickness for each eye from a single mouse).

**Table 1.** Retinal thicknesses with various promoters. Mean retinal thickness measured from the nerve fiber layer (NFL) to the retinal pigment epithelium (RPE) or the outer plexiform layer (OPL) to the RPE at 2 months and 4 months of age. Measurements represent the thickness in the inferior region of the retina (closest to the injection site). All treated eyes were treated with scAAV9 with the respective promoter driving expression of hNMNAT1. The virus was delivered at 1E9 vg/μl and co-injected with AAV9 CMV.GFP at a titer of 1E8 vg/μl.

| Table 1 | | Mean NFL to RPE<br>Thickness (μm) | | Mean OPL to RPE<br>Thickness (μm) | |
| --- | --- | --- | --- | --- | --- |
| Promoter | Genotype | 2 Months | 4 Months | 2 Months | 4 Months |
| CASI | WT | 222.46 | 210.75 | 125.42 | 119.04 |
|  | *Nnmnat1*<sup>V9M/V9M</sup> | 215.95 | 191.6 | 119.1 | 106.45 |
| hGRK1 | WT | 222.53 | 217.27 | 125.69 | 122.97 |
|  | *Nnmnat1*<sup>V9M/V9M</sup> | 201.03 | 182.89 | 102.93 | 93.5 |
| ProC1 | WT | 222.92 | 217.56 | 128.06 | 124.72 |
|  | *Nnmnat1*<sup>V9M/V9M</sup> | 203.88 | 182.56 | 109.03 | 95.34 |
| VMD2 | WT | 214.25 | 210.67 | 124.08 | 123.25 |
|  | *Nnmnat1*<sup>V9M/V9M</sup> | 163.25 | 146.34 | 71.7 | 58.9 |
| ProC3 | WT | 221.79 | 219.9 | 124.29 | 124.81 |
|  | *Nnmnat1*<sup>V9M/V9M</sup> | 194.25 | 180.6 | 101.5 | 93.34 |
| ProD5 | WT | 222.36 | 217.36 | 126.32 | 124.07 |
|  | *Nnmnat1*<sup>V9M/V9M</sup> | 201.2 | 185.41 | 108.83 | 101.88 |
| Untreated | WT | 226.61 | 221.99 | 130.47 | 128.85 |
|  | *Nnmnat1*<sup>V9M/V9M</sup> | 154.8 | 125.25 | 61.66 | 39.09 |

To better address whether expression of NMNAT1 solely in the photoreceptors is sufficient to prevent retinal degeneration in NMNAT1-associated disease, we tested the ProC1 promoter, a synthetic promoter that has been reported to drive gene expression in both rod and cone photoreceptor cells (15). We confirmed that delivery of scAAV9.ProC1.hNMNAT1.WPRE resulted in expression of hNMNAT1 in both the rod and cone photoreceptor cells, but not other retinal cell types (Figure 1C). Interestingly, we found that expression solely in the photoreceptor cells in the retina was sufficient to prevent retinal degeneration, preserving retinal thickness at both 2 and 4 months of age (Figure 2C, Figure S3C). The level of preservation was similar to the of treatment with scAAV9.hGRK1.hNMNAT1.WPRE and was slightly lower than the preservation with treatment with scAAV9.CASI.hNMNAT1.WPRE, although the differences were not statistically significant (Figure 3, Table 1).

We next aimed to assess whether NMNAT1 expression in the retinal pigment epithelium (RPE) is sufficient to prevent retinal degeneration, which would indicate that the RPE are involved in *NMNAT1*-associated disease pathology. We delivered scAAV9.VMD2.hNMNAT1.WPRE to express hNMNAT1 specifically in the RPE cells of the retina. IHC shows expression of hNMNAT1 in the RPE cells, with no expression in the neural retina (Figure 1D). Expression of hNMNAT1 in the RPE was not sufficient to prevent loss of retinal cells. There were minimal differences in total retinal thickness (Figure S3D) and photoreceptor layer thickness (Figure 2D) at both 2 and 4 months of age in *Nmnat1^V9M/V9M^* retinas treated with scAAV9.VMD2.hNMNAT1.WPRE as compared to untreated retinas. Interestingly, there was a small benefit observed in the superior region of treated retinas at 4 months of age (Figure 3, Figure S3H), suggesting that NMNAT1 may play a minor role in the RPE function.

Although we found that expression of hNMNAT1 in the photoreceptor cells was sufficient to prevent retinal degeneration, we aimed to assess whether expression of hNMNAT1 in other neural retinal cell types, not including the photoreceptor cells, could confer a benefit. To this end, we treated mice with scAAV9.ProC3.hNMNAT1.WPRE, using the ProC3 synthetic promoter reported to drive expression of transgenes in the inner retina. At 2- and 4-weeks post-injection, we observed hNMNAT1 expression in the retinal ganglion cell layer, with some leakiness in the photoreceptor layer (Figure 1E). Surprisingly, this expression pattern was sufficient to provide some benefit against retinal degeneration in *Nmnat1^V9M/V9M^* mice (Figure 2E, Figure S3E). However, at two months of age, the retinal thickness in the scAAV9.CASI.hNMNAT1.WPRE-treated retinas was significantly greater than those of the scAAV9.ProC3.hNMNAT1.WPRE-treated retinas (Figure 3, Table 1), suggesting that expression of NMNAT1 in other cell types or expression in the photoreceptors at a greater level is essential for proper retinal function.

We identified that other than expressing NMNAT1 in most retinal cell types using the CASI promoter, the greatest benefit was observed using promoters that led to expression in the photoreceptor cells. Therefore, we aimed to determine whether expression of NMNAT1 is required in both rod and cone photoreceptor cells, or only one cell type. To assess whether expression of NMNAT1 in rods is sufficient to prevent retinal degeneration, we treated mice with scAAV9.ProD5.hNMNAT1.WPRE. ProD5 is a synthetic promoter reported to drive expression specifically in rod photoreceptor cells (15). We confirmed expression in the rod photoreceptor cells (Figure 1F), and lack of hNMNAT1 staining the cone photoreceptor cells with co-staining for the cone marker, peanut agglutinin (PNA) (data not shown). Expression of hNMNAT1 in the rods led to significant preservation of retinal structure in treated *Nmnat1^V9M/V9M^* mice as compared to the untreated contralateral eye (Figure 2F, Figure S3F). The level of preservation was similar to that of other promoters that expressed in both rods and cones (ProC1 and hGRK1) (Figure 3, Table 1). These results suggest that NMNAT1 plays an essential role in rod photoreceptor cells.

We next aimed to assess the role of NMNAT1 in the cone photoreceptor cells. To this end, the mouse opsin (mOPs) promoter was used to drive NMNAT1 expression in rod photoreceptor cells (17). Mice were treated with scAAV9.mOPs.hNMNAT1.WPRE. Unfortunately, in our hands, this promoter was not specific to the cone photoreceptor cells and led to expression also in the rod photoreceptor cells as seen by the localization of the hNMNAT1 expression, (Figure S4A), and by PNA staining to label cones (data not shown). As this promoter was not cone-specific, assessment of structural rescue via OCT was not carried out. We next utilized a promoter that is a combination of the enhancer from the *IRBP* gene and the promoter from the *GNAT2* gene, which has been shown previously to express specifically in the cone photoreceptor cells (18). Although we did observe some staining in the cones with this promoter, it was very minimal (Figure S4B). Protection was not observed at 2 months of age, although some protection was observed at 4 months of age (Figure S4C-D). This protection was significantly lower than that observed with any of the other promoters, besides VMD2, at both 2 and 4 months of age. These findings suggest that expression of NMNAT1 in the cones alone is not sufficient to prevent retinal degeneration and that rods also require NMNAT1 expression at least in this model of *NMNAT1*-associated disease, where rapid photoreceptor degeneration occurs.

Since we were unable to detect strong expression of hNMNAT1 specifically in cone photoreceptor cells using putative cone-specific promoters, we wanted to assess cone viability in *Nmnat1^V9M/V9M^* mice without treatment with our AAV constructs. We aimed to gain a better understanding of whether there are differences in the localization or number of cones in *Nmnat1^V9M/V9M^* retinas vs. WT retinas at the same age. To assess cones, we stained retinal sections from *Nmnat1^V9M/V9M^* retinas vs. WT mice prior to when loss of retinal thickness is observed in the *Nmnat1^V9M/V9M^* strain (2 and 3 weeks of age) and when loss of cells is apparent as measured by thinning of the photoreceptor layer (4 weeks of age) (9). Sections were stained with the cone marker PNA and the number of cones per area were manually quantified. Interestingly, although a consistent number of cones were observed in WT mice at 2, 3, and 4 weeks of age, *Nmnat1^V9M/V9M^* mice exhibited loss of cone photoreceptor cells starting at 3 weeks of age (Figure 4 A-B), prior to when measurable thinning of the retina is observed. Cones were further depleted in *Nmnat1^V9M/V9M^* mice by 4 weeks of age (Figure 4 A-B). These findings suggest that cone photoreceptor cells are depleted first in the *Nmnat1^V9M/V9M^* retina, indicating that cones may be uniquely affected by loss of NMNAT1. We also aimed to assess whether cones were preserved long-term in the treated *Nmnat1^V9M/V9M^* mice. We stained sections from 9 month old mice treated with scAAV9.CASI.hNMNAT1.WPRE with the PNA cone marker. Although the photoreceptor cells are not present in the untreated eye, we see preservation of cone photoreceptors in the treated *Nmnat1^V9M/V9M^* retina at 9 months of age (Figure 4C). These findings demonstrate that long-term preservation of the cones is achievable with NMNAT1 expression throughout the retina.

**Figure 4.**
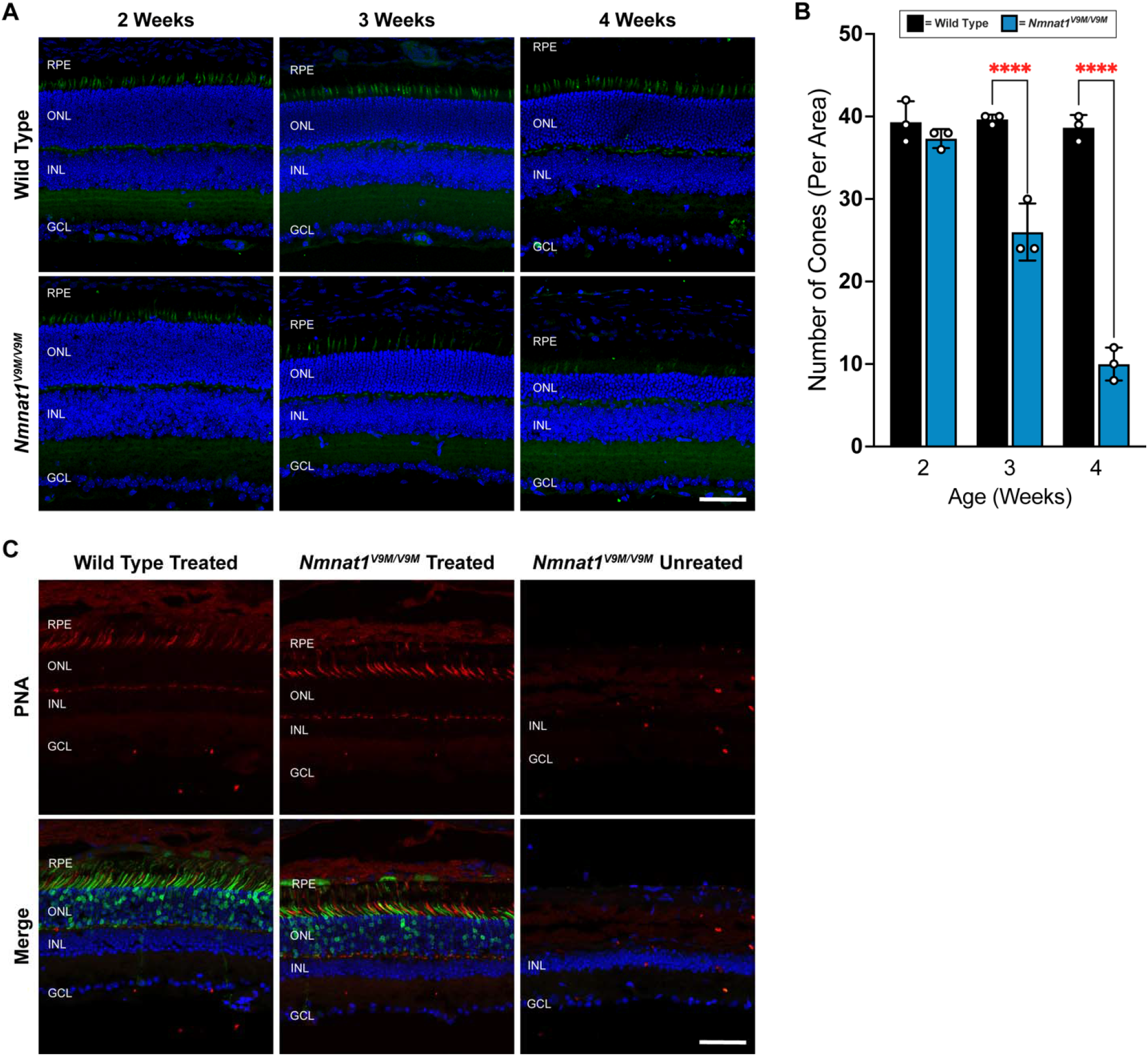
Cone degeneration occurs before measurable loss of retinal thickness. (A) Expression of PNA (a cone-specific marker, green) in wild type and untreated *Nmnat1^V9M/V9M^* mice at 2, 3, and 4 weeks of age. Bottom panels are counterstained with DAPI (blue). Abbreviations: retinal pigment epithelium (RPE); outer nuclear layer (ONL); inner nuclear layer (INL); ganglion cell layer (GCL). Scale bar indicates 20 μm. (B) Quantification of the number of cones per area based on PNA staining. Statistically significant differences are noted by red asterisks (n = 3 WT and n = 3 *Nmnat1^V9M/V9M^* mice for each age, 2Way ANOVA, multiple comparisons, *P* < 0.05, error bars represent the SEM, points represent the number of cones per image from a single mouse). (C) Expression of PNA (red) in wild type (left panel) and *Nmnat1^V9M/V9M^* retinas (middle panel) co-injected with scAAV9.CASI.hNMNAT1.WPRE and scAAV2/9.CMV.EGFP or untreated (right panel) at 9 months of age. Bottom panels are counterstained with DAPI (blue). Abbreviations: retinal pigment epithelium (RPE); outer nuclear layer (ONL); inner nuclear layer (INL); ganglion cell layer (GCL). Scale bar indicates 20 μm.

Finally, we confirmed that hNMNAT1 expression with each promoter did not lead to retinal toxicity or damage. The inferior and superior regions of WT littermate mice treated with the various constructs did not display retinal thinning as compared to the contralateral untreated WT eye, at either 2 or 4 months of age, with any of the AAVs that were tested (Figures S5-S6).

## Discussion

In the present study, we aimed to determine which cell types in the retina require NMNAT1 expression and to gain a better understanding of retinal cell types that should be targeted in a gene therapy for *NMNAT1*-associated disease and to enhance our understanding of disease biology. To this end, we systematically expressed hNMNAT1 using AAV and cell type-specific promoters to drive the expression of hNMNAT1 in the retinas of *Nmnat1^V9M/V9M^* mice. The data show that expression of hNMANT1 in the photoreceptors is sufficient to prevent significant retinal degeneration in this model. Expression of NMNAT1 in the RPE does not significantly prevent retinal degeneration in *Nmnat1^V9M/V9M^* mice. We also show that expression of hNMNAT1 in the rod photoreceptors alone is sufficient to prevent retinal degeneration. Finally, we demonstrate that cone photoreceptor cells are reduced in *Nmnat1^V9M/V9M^* mice, prior to a measurable loss of retinal thickness. These findings suggest that NMNAT1 expression is required in photoreceptors, that cone photoreceptors are uniquely affected in disease, and that NMNAT1 likely plays an important role in other neural retina cell types.

Expression of hNMNAT1 in most retinal cell types, driven by the CASI promoter, provided the greatest preservation of retinal thickness. However, expression of NMNAT1 with promoters that led to expression primarily in the photoreceptors also led to significant preservation of the retina. Consistent with our findings, a previous study has shown that expression of NMNAT1 specifically in the photoreceptor cells, with delivery of AAV encoding NMNAT1 with expression driven by the hGRK1 promoter, is sufficient to prevent retinal degeneration with conditional knockout of NMNAT1 in the retina (19). These findings suggest that expression of NMNAT1 in the photoreceptors alone can confer a significant benefit in NMNAT1-associated disease, and that photoreceptors likely play a significant role in disease pathology. These data are consistent with previous findings from our group regarding the molecular mechanisms underlying NMNAT1-associated disease (12). Other evidence also suggests that photoreceptors are exceptionally vulnerable to loss of NMNAT1. Knockout of *Nmnat1* specifically in the retina at embryonic day 9.5-12.5 results in developmental defects and rapid retinal degeneration (20). Early embryonic knockout of NMNAT1 indicate that it plays an essential role in terminal differentiation of photoreceptor cells and that this phenotype is driven by dysregulation of transcription in photoreceptor cells due to loss of NMNAT1 expression (20). These findings indicate that at least some expression of NMNAT1 is required for proper retinal development.

Mouse models with ubiquitous conditional knockout of *Nmnat1* also exhibit retinal degeneration (19). Cell type-specific knockout of *Nmnat1* showed that NMNAT1 expression was required in both the rod and cone photoreceptors (19). Knockout of *Nmnat1* specifically in the cone photoreceptor cells did not result in noticeable histological differences, but there was complete loss of the photopic b-wave on electroretinogram (ERG) testing, a measurement of cone function (19). Knockout of *Nmnat1* in rod photoreceptors led to a similar level of degeneration as ubiquitous knockout of *Nmnat1* (19), suggesting that the photoreceptors are primarily contributing to the retinal degeneration phenotype in *NMNAT1*-associated disease. Together, these results support the hypothesis that the photoreceptors are uniquely vulnerable to loss of NMNAT1 function, that NMNAT1 expression is essential for the proper function of both rod and cone photoreceptor cells, and that NMNAT1 expression in photoreceptor cells alone is sufficient to prevent retinal degeneration.

In the present study, we also aimed to assess whether expression of hNMNAT1 in the cone photoreceptor cells alone was sufficient to prevent retinal degeneration. We were unable to successfully target only the cone photoreceptors using the mOPs promoter (Figure S4). When we used the cone specific IRBPeGNAT2p promoter we observed only sparce expression of hNMNAT1 in cones, with very minimal preservation of retinal thickness in treated *Nmnat1^V9M/V9M^* mice as compared to untreated mice (Figure S4). We hypothesize that the differences in cell type-specific expression observed in our hands may be due to elements in our constructs that may affect promoter activity or differences due to the nature of scAAV, as recent evidence suggests that there is intrinsic promoter activity in the AAV ITRs themselves (21). Interestingly, when staining retinal sections with the cone-specific marker PNA, we observed a reduction in the number of cones in *Nmnat1^V9M/V9M^* retinas at 3 weeks of age, which is prior to when a measurable difference in retinal thickness is observed in this model (9). The loss of cones continues at 4 weeks of age, when retinal thinning is observed. Although we do not detect a significant portion of rod loss at 3 weeks of age, as no retinal thinning is yet observed, we have previously reported that we observe TUNEL-positive rod cells at 4.5 weeks and 6 weeks of age, with significant retinal thinning beginning at 4 weeks of age (9, 11). These findings suggest that the cones are the first cells to degenerate in the retinas of *Nmnat1^V9M/V9M^* mice, followed by the loss of rods.

Consistent with this hypothesis, patients with mutations in NMNAT1 exhibit macular atrophy (1, 5). Interestingly, evidence suggests that cone photoreceptors have greater energy requirements than rod photoreceptors (22), which may explain why cones are uniquely affected in *NMNAT1*-associated disease. We hypothesize that we are unable to prevent retinal degeneration and efficiently express hNMNAT1 solely in the cone photoreceptor cells in *Nmnat1^V9M/V9M^* mice because these cells are already depleted by 3 weeks of age, so treatment at 2 weeks of ages does not provide a sufficient time window for NNMNAT1 to be expressed and prevent the degeneration of cones. Additionally, studies indicate that rods are required for proper cone function (23). We hypothesize that expression of NMNAT1 is required in the rods, as its expression in the cones is not sufficient to prevent their degeneration.

Together, our findings suggest that NMNAT1 plays an essential role in retinal photoreceptor cells. Expression of NMNAT1 in photoreceptor cells alone is sufficient to prevent retinal degeneration, suggesting that a gene replacement therapy for *NMNAT1*-associated disease must target these cells. Future studies using single cell RNAseq will aim to develop a better understanding of the cell-type specific effects of mutations in *NMNAT1*. Other models, such as retinal organoids derived from human iPSCs with the p.V9M mutation in *NMNAT1* will also provide more insight about the cell type-specific role of NMNAT1 and the specific roles of NMNAT1 in rod vs. cone photoreceptor cells.

## Materials and Methods

### Mouse Lines

The *Nmnat1^V9M/V9M^* mouse line was derived and outcrossed as described previously (9). *Nmnat1^V9M/WT^* mice were bred together, so that *Nmnat1^V9M/V9M^* and *Nmnat1^WT/WT^* mice from the same litter could be used for experiments. Male and female mice were used without preference. Mice were given a 4% fat rodent diet and water *ad libitum* and housed with a 12-hour light/12-hour dark cycle. All procedures were approved by the Animal Care and Use Committee of Schepen’s Research Institute of Mass Eye and Ear and conformed to the Association for Research in Vision and Ophthalmology Statement for the Use of Animals in Ophthalmic and Vision Research.

### Genotyping

Genotyping for *Nmnat1* p.V9M was performed as described previously (9). In brief, PCR was performed on DNA extracted from a tail biopsy sample using the forward primer 5’-CATGGCTGTGCTGAGGTG-3’ (intron 1) and reverse primer 5’-AACAGCCTGAGGTGCATGTT-3’ (exon 2). These primers amplify a 691 bp region of *Nmnat1*. Sanger sequencing using the primer 5’-ACGTATTTGCCCACCTGTCT-3’ was used to identify the genotypes of the mice at position c.25 as being *Nmnat1^WT/WT^, Nmnat1^WT/V9M^*, or *Nmnat1^V9M/V9M^*.

### DNA Construct Design and Preparation

DNA 2.0 (Menlo Park, CA) was used to design codon optimized human *NMNAT1* cDNA, which was synthesized into a gBlock gene fragment (Integrated DNA Technologies, Coralville, IA). The hNMNAT1 gBlock was inserted into a self-complimentary AAV plasmid. Cell type specific promoter sequences were inserted into this base plasmid. Synthetic cell type specific promoters were obtained from the following Addgene plasmids: 161_pAAV-ProD5-CatCh-GFP-WPRE (Addgene plasmid # 125981; http://n2t.net/addgene:125981; RRID:Addgene_125981), 124_pAAV-ProC3-CatCh-GFP-WPRE (Addgene plasmid # 125939; http://n2t.net/addgene:125939; RRID:Addgene_125939), and 123_pAAV-ProC1-CatCh-GFP-WPRE (Addgene plasmid # 125937; http://n2t.net/addgene:125937; RRID:Addgene_125937) which were gifts from Botond Roska (15). The VMD2 and hGRK1 promoters were synthesized into gBlock gene fragments (Integrated DNA Technologies, Coralville, IA) and inserted into the base a self-complimentary AAV plasmid. The CASI promoter construct was developed as described previously (14). All promoter sequences can be found in Table S1.

### AAV Preparation

Plasmids with the desired constructs were generated using an endotoxin free maxiprep kit (12362, QIAGEN, Germantown, MD) and confirmed via whole plasmid NGS. The AAV preparations were made by the Grousbeck Gene Therapy Center of Massachusetts Eye and Ear, as described previously (24). The final purified virus was titered and diluted in buffer containing 1X PBS, 35mM NaCl, and 0.001% Pluronic F68 surfactant (24040032, ThermoFisher, Waltham, MA). For injections, < 0.25% of fluorescein (NDC: 17478-253-10, AK-Fluor, Akorn, Lake Forest, IL) was added to allow visualization of the injection fluid.

### Subretinal Injections

All mice were injected with the respective AAV vectors at 2 weeks of age (P14). To anesthetize the mice, a mixture of 37.5 mg/kg ketamine and 3.8 mg/kg xylazine was administered via intraperitoneal (IP) injection. Proparacaine HCl (0.5%) drops were placed on the cornea to provide local anesthesia and the pupils were dilated with a mixture of tropicamide (0.25%) and phenylephrine HCl (1.25%). The eye was then proptosed and a 30G needle was used to pierce the superior portion of the eye at the corneal scleral divide, visualized using a dissection scope. A 33G blunt cannula attached to the Micro4 microinjection pump with RPE kit (World Precision Instruments, Sarasota, FL) was inserted into the eye, and positioned to deliver 0.75 μL of the vector into the subretinal space in the inferior-nasal region of the eye. The formation of a bleb was confirmed visually, and the cannula tip was held in place for about 3 seconds after the injection to prevent leakage. After removal of the cannula, a sterile eye spear (400101, BVI Medical, Waltham, MA) was held at the injection site for several seconds. The eyes were then treated with lubricant gel (Genteal, Alcon, Geneva, Switzerland) and mice were given 2mg/kg Yohimbine HCl (Wedgewood Pharmacy, Swedesboro, NJ) delivered via subcutaneous injection to prevent the development of corneal opacities (25, 26). Mice were placed on a heating pad to recover from the anesthesia.

### Optical Coherence Tomography (OCT) and Fundoscopy

Mice were anesthetized with a mixture of 37.5 mg/kg ketamine and 3.8 mg/kg xylazine delivered via IP injection. The pupils were dilated with a mixture of tropicamide (0.25%) and phenylephrine HCl (1.25%). Spectral-domain OCT and InVivoVue OCT software (Bioptogen) was used to obtain cross-sectional images of the retina and quantifications of the retinal thickness, as described previously (14). To confirm success of the injection, fundus photography with a filter for EGFP was used to visualize EGFP expression in the retina.

### Immunohistochemistry (IHC)

Tissue collection and IHC were performed as described previously (12). In brief, mice were euthanized via CO_2_ asphyxiation and the eye was enucleated. Eyes were washed in PBS and then soaked in 4% paraformaldehyde (PFA) (v/v) for 4 minutes. The cornea and lens were then removed, and the posterior cup was incubated in 4% PFA for 20 minutes. After washing in PBS, eyecups were stored in 30% sucrose (w/v) overnight. After embedding in optimal cutting temperature (OCT) medium, eyes were sectioned to 10 μm thickness. Sections were stained with an antibody specific to hNMNAT1, described previously (14). The slides were imaged using the TCS SP8 confocal microscope (Leica, Wetzlar, Germany). Image quantification of cones was performed by a masked individual. The number of cones per total area of each image was manually quantified from areas of the retina equidistant from the optic nerve.

### Statistics

Prism (Version 9, GraphPad, San Diego, CA) was used to produce graphs and perform statistical analysis. 2Way ANOVA with multiple comparisons was used to assess differences between groups. Data are presented as the mean ± S.E.M and a *P*-value of < 0.05 was considered statistically significant.

## Acknowledgements

This work was supported by the National Eye Institute [EY012910 (E.A.P) and P30EY003790 (Core Grant)].

## Author Contributions

E.E.B. designed research studies, conducted experiments, acquired data, analyzed data, and wrote the manuscript. M.J.S. designed research studies, conducted experiments, acquired data, analyzed data, and contributed to writing the manuscript.

E.A.P. designed research studies and edited the manuscript.

## Data Availability Statement

The datasets generated during and/or analyzed during the current study are available from the corresponding author on reasonable request.

