## Supplemental Materials for "Expression of NMNAT1 in the Photoreceptors is Sufficient to Prevent *NMNAT1*-Associated Disease"

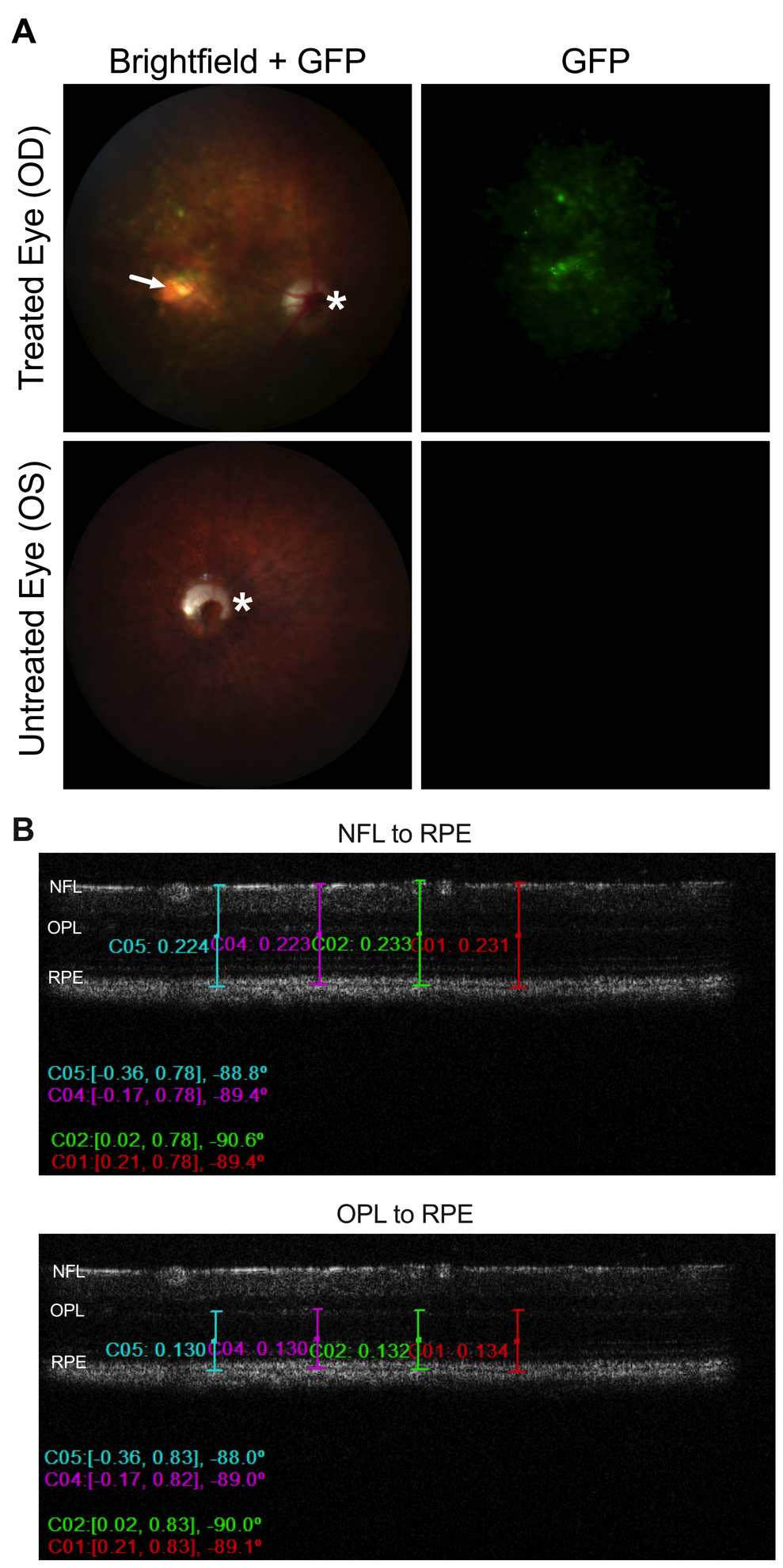

**Figure S1. Fundus photography and optical coherence tomography.** (A) Fundus images showing a representative treated eye (top row) and untreated eye (bottom row) in the brightfield and GFP filter channels. * Indicates the optic nerve head. The arrow indicates the injection site. (B) Representative OCT images showing how the measurements of retina thickness were measured from the nerve fiber layer (NFL) to retinal pigment epithelium (RPE) (top image) or from the outer plexiform layer (OPL) to the RPE (bottom image).

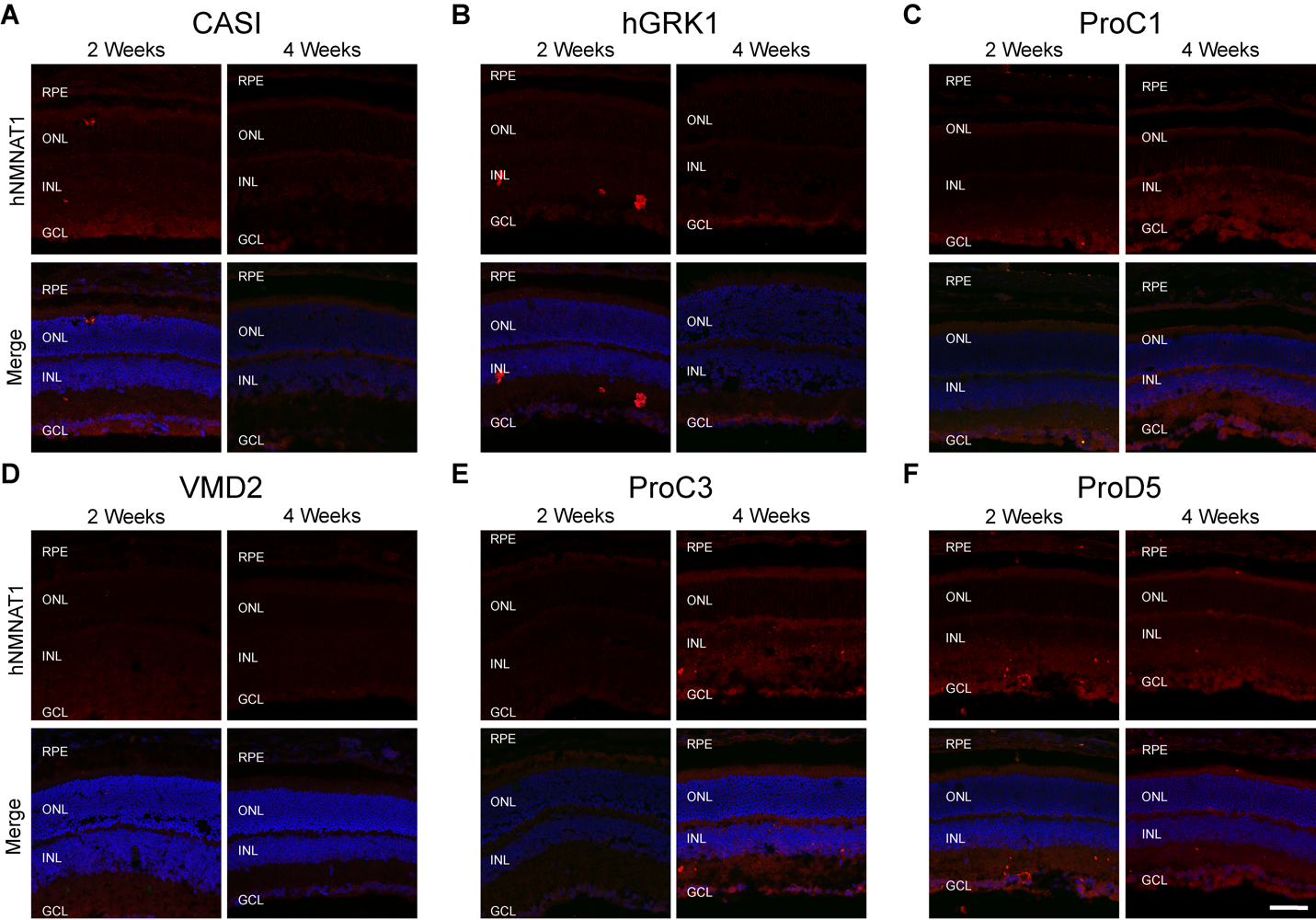

**Figure S2. hNMNAT1 expression is not present in the untreated contralateral eye.** Immunohistochemistry from the contralateral untreated eye. Top panels show lack of staining with an antibody specific to human NMNAT1 (hNMNAT1) (red) two weeks and 4 weeks post injection in the contralateral eye. Any red signal is autofluorescence. Bottom panels show counterstaining with DAPI (blue) and show no signal in the green channel (no GFP signal). The treatments administered to the contralateral eyes include delivery of AAV9 with hNMNAT1 expression driven by the (A) CASI promoter, (B) ProC1 promoter, (C) VMD2 promoter, (D) ProC3 promoter, (E) ProD5 promoter, and (F) hGRK1 promoter. Note that any fluorescence seen is autofluorescence. Abbreviations: retinal pigment epithelium (RPE); outer nuclear layer (ONL); inner nuclear layer (INL); ganglion cell layer (GCL). Scale bars indicate 20 µm.

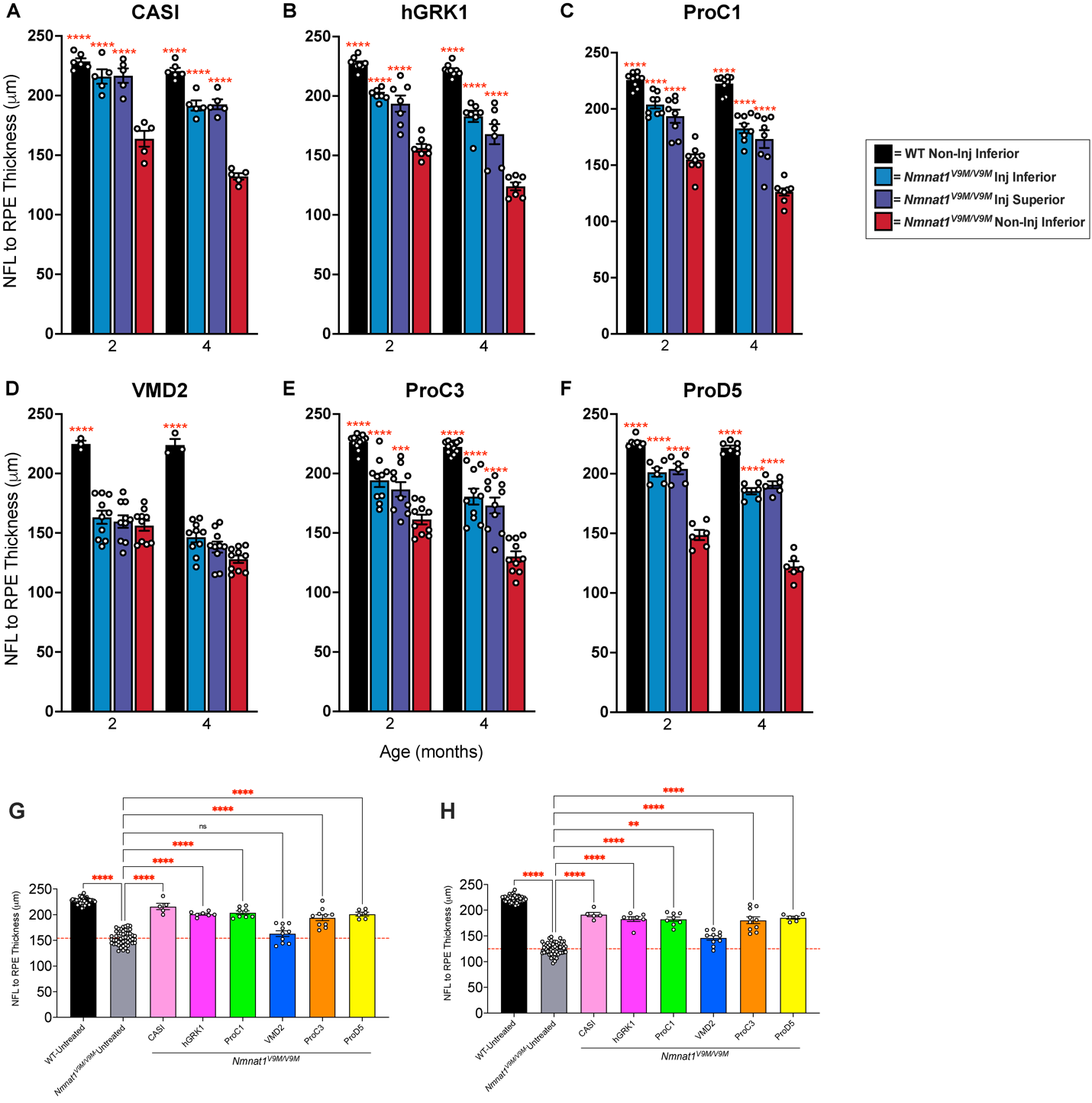

**Figure S3.** **Retinal thickness is preserved with expression of NMNAT1 in the neural retina.** Measurements of retinal thickness measured from the nerve fiber layer (NFL) to the retinal pigment epithelium (RPE). Measurements are based on SD-OCT imaging of retinas from mice treated with scAAV9 with promoters driving expression of hNMNAT1 in various cell types. (A-F) Measurements include the retinal thickness of untreated WT mice (black), the inferior region of treated *Nmnat1^V9M/V9M^* retinas, adjacent to the injection site (blue), the superior region of treated *Nmnat1^V9M/V9M^* retinas, distal to the injection site (purple), and untreated *Nmnat1^V9M/V9M^* contralateral eyes (red). The promoters used include (A) CASI (most cell types), (B) hGRK1 (rods and RPE), (C) ProC1 (photoreceptors), (D) VMD2 (RPE), (E) ProC3 (ganglion cells), (F) ProD5 (rods). Measurements of retinal thickness measured from the nerve fiber layer (NFL) to RPE at 2 months (G) and 4 months of age (H). Measurements for *Nmnat1^V9M/V9M^* untreated retinas represent the average retinal thickness for the untreated contralateral eye for all groups. Wild type (WT) untreated measurements represent the average retinal thickness for the untreated contralateral eye from all WT groups. Measurements represent the thickness in the inferior region of the retina (closest to the injection site). All treated eyes were treated with scAAV9 with the respective promoter driving expression of hNMNAT1. This virus was delivered at 1E9 vg/μl and co-injected with AAV9 CMV.GFP at a titer of 1E8 vg/μl. The red dashed line represents the mean thickness of the untreated *Nmnat1^V9M/V9M^* retinas at the respective age. Statistically significant preservation of retinal thickness between untreated and treated *Nmnat1^V9M/V9M^* retinas are noted by red asterisks (for all graphs: n $\geq$ 3 WT and n $\geq$ 5 *Nmnat1^V9M/V9M^* mice for each group at each age, 2Way ANOVA, multiple comparisons, *P* < 0.05, error bars represent the SEM, points represent the respective retinal thickness for each eye from a single mouse). Abbreviations: nerve fiber layer (NFL); retinal pigment epithelium (RPE); inner nuclear layer (INL); ganglion cell layer (GCL); spectral-domain optical coherence tomography (SD-OCT).

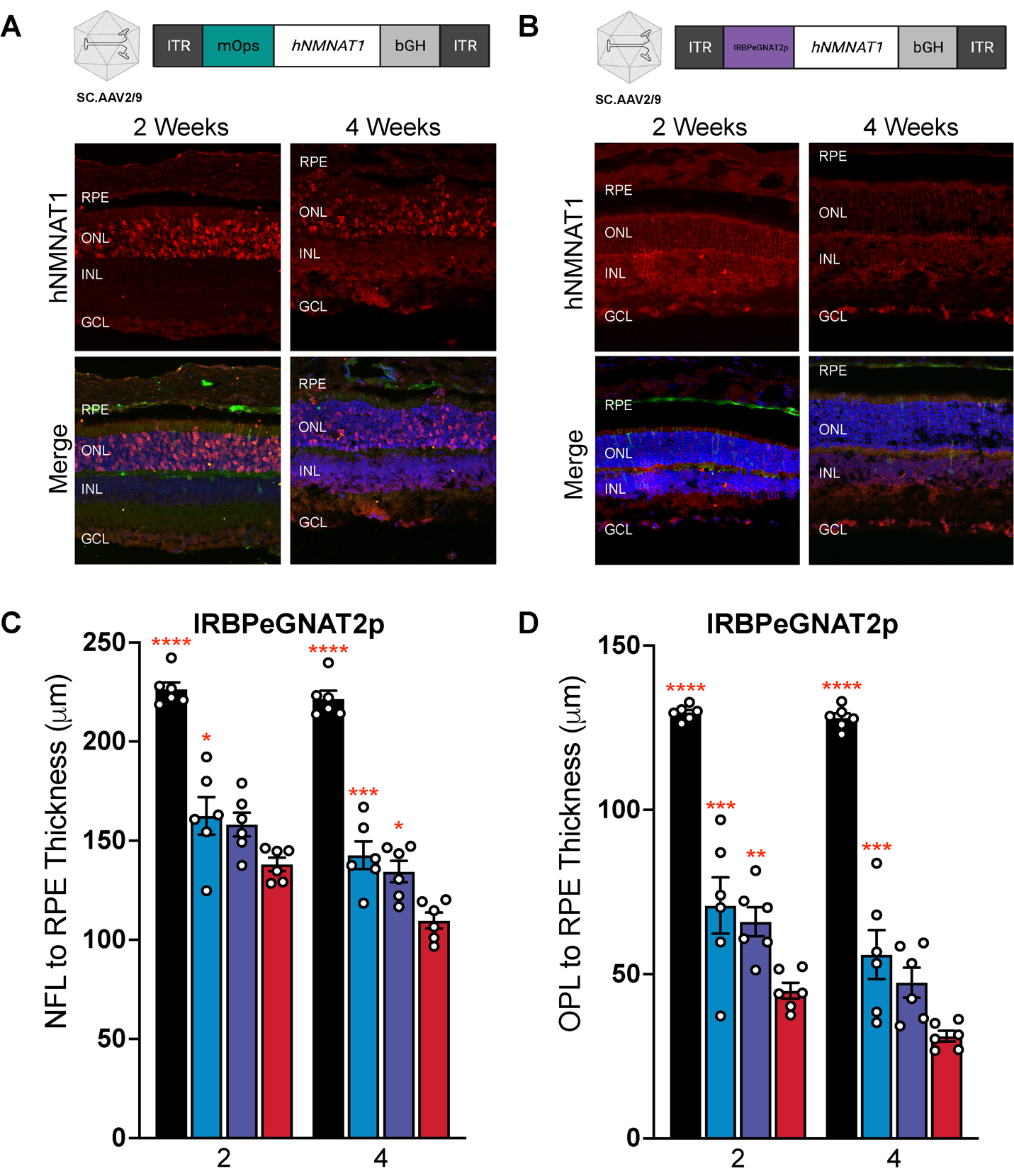

**Figure S4. Expression of NMNAT1 in cone photoreceptor cells.** (A-B) Diagrams show the design of AAV constructs with various promoters. Immunofluorescence images demonstrate the cell type-specific expression of hNMNAT1 (red) after delivery scAAV9 at a titer of 1E9 vg/μl, 2- and 4-weeks post subretinal injection. Expression of NMNAT1 using (A) the mOPs promoter, which led to expression in photoreceptor cells and (B) IRBPeGNAT2p which led to minimal expression in cone photoreceptor cells. Abbreviations: retinal pigment epithelium (RPE); outer nuclear layer (ONL); inner nuclear layer (INL); ganglion cell layer (GCL). Top panels show staining with an antibody specific to human NMNAT1 (hNMNAT1). Bottom panels are counterstained with DAPI (blue). Scale bar indicates 20 µm. (C-D) Measurements of retinal thickness from mice treated with AAV2/9.IRBP2GNAT2p.hNMNAT1.WRPE measured from the nerve fiber layer (NFL) to the retinal pigment epithelium (RPE) (C) or the outer plexiform layer (OPL) to the RPE (D). Measurements include the retinal thickness of untreated WT mice (black), the inferior region of treated *Nmnat1^V9M/V9M^* retinas, adjacent to the injection site (blue), the superior region of treated *Nmnat1^V9M/V9M^* retinas, distal to the injection site (purple), and untreated *Nmnat1^V9M/V9M^* contralateral eyes (red).

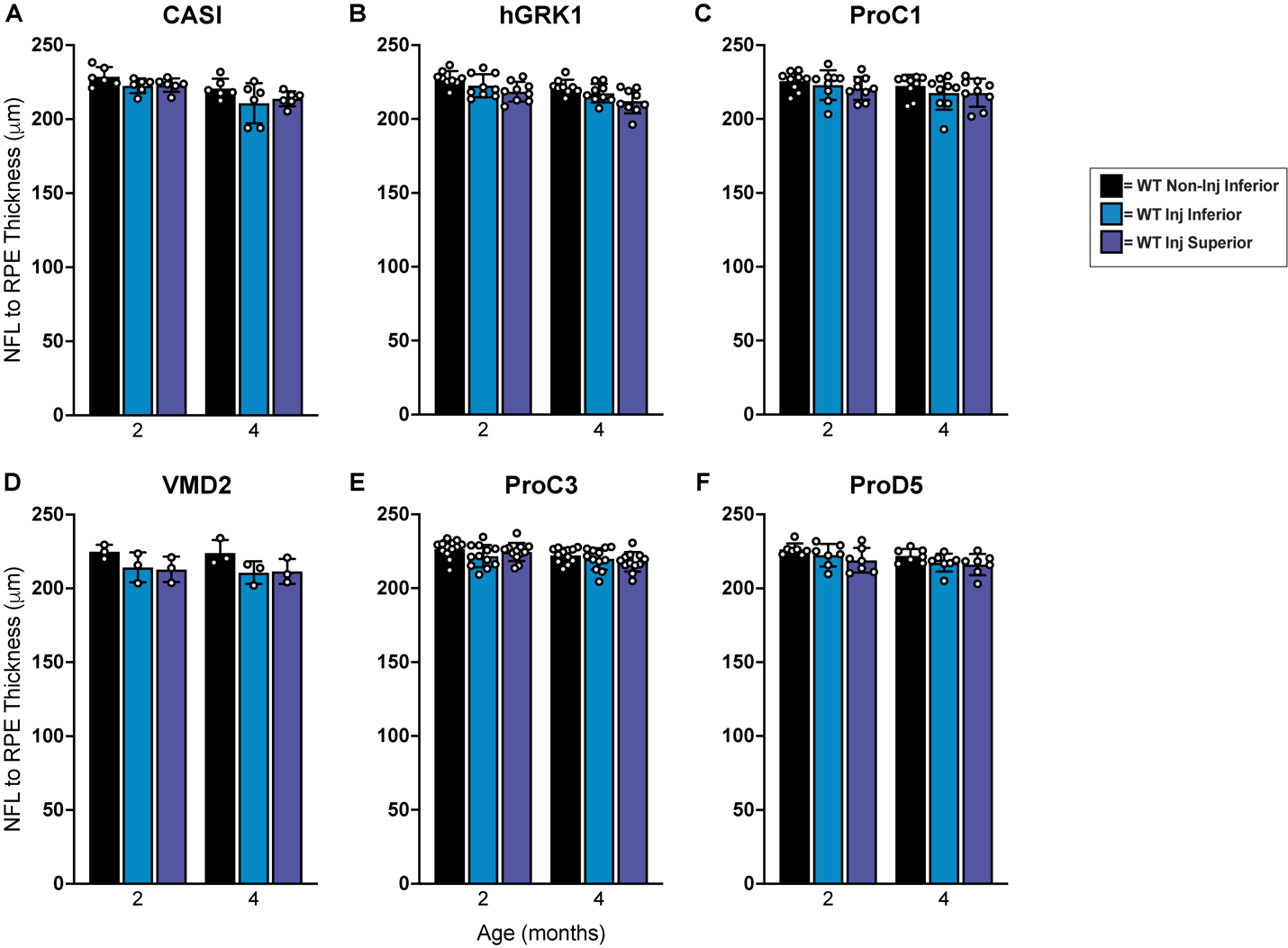

**Figure S5. Total retinal thickness measurements show expression of hNMNAT1 is not toxic in the WT retina.** Measurements of retinal thickness measured from the nerve fiber layer (NFL) to the retinal pigment epithelium (RPE). Measurements are based on SD-OCT imaging of retinas from mice treated with scAAV9 with promoters driving expression of hNMNAT1 in various cell types. Measurements include the retinal thickness of untreated WT mice (black), the inferior region of treated WT retinas, adjacent to the injection site (blue), and the superior region of treated WT retinas, distal to the injection site (purple). The promoters used include (A) CASI (most cell types), (B) ProC1 (photoreceptors), (C) VMD2 (RPE), (D) ProC3 (ganglion cells), (E) ProD5 (rods), (F) hGRK1 (rods and some RPE). There are no significant differences in the retinal thickness between treated and untreated retinas (n $\geq$ 3 mice per group, 2Way ANOVA, multiple comparisons, *P* > 0.05, error bars represent the SEM, points represent the respective retinal thickness for each eye from a single mouse).

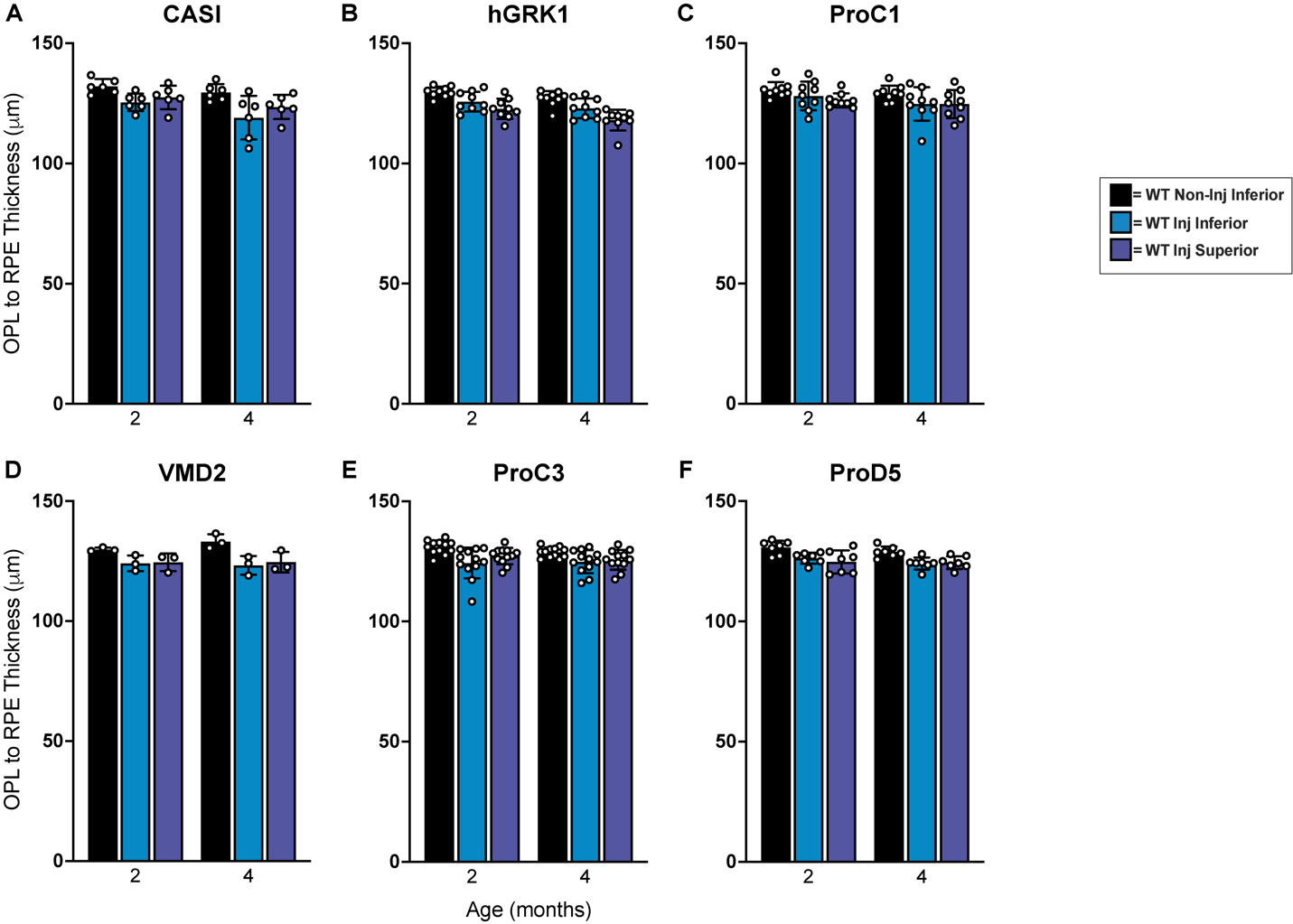

**Figure S6. Photoreceptor thickness measurements show expression of hNMNAT1 is not toxic in the WT retina.** Measurements of retinal thickness measured from the outer plexiform layer (OPL) to the retinal pigment epithelium (RPE). Measurements are based on SD-OCT imaging of retinas from mice treated with scAAV9 with promoters driving expression of hNMNAT1 in various cell types. Measurements include the retinal thickness of untreated WT mice (black), the inferior region of treated WT retinas, adjacent to the injection site (blue), and the superior region of treated WT retinas, distal to the injection site (purple). The promoters used include (A) CASI (most cell types), (B) ProC1 (photoreceptors), (C) VMD2 (RPE), (D) ProC3 (ganglion cells), (E) ProD5 (rods), (F) hGRK1 (rods and some RPE). There are no significant differences in the retinal thickness between treated and untreated retinas (n $\geq$ 3 mice per group, 2Way ANOVA, multiple comparisons, *P* > 0.05, error bars represent the SEM, points represent the respective retinal thickness for each eye from a single mouse).

| **Table S1**  Promoter name, sequence used, cell type expression, and references, if applicable. | | | |
| --- | --- | --- | --- |
| Promoter | Promoter Sequence | Cell Expression | Reference |
| CASI | ggagttccgcgttacataacttacggtaaatggcccgcctggctgaccgcccaacgacccccgcccattgacgtcaataatgacgtatgttcccatagtaacgccaatagggactttccattgacgtcaatgggtggagtatttacggtaaactgcccacttggcagtacatcaagtgtatcatatgccaagtacgccccctattgacgtcaatgacggtaaatggcccgcctggcattatgcccagtacatgaccttatgggactttcctacttggcagtacatctacgtattagtcatcgctattaccatggtcgaggtgagccccacgttctgcttcactctccccatctcccccccctccccacccccaattttgtatttatttattttttaattattttgtgcagcgatgggggcggggggggggggggggcgcgcgccaggcggggcggggcggggcgaggggcggggcggggcgaggcggagaggtgcggcggcagccaatcagagcggcgcgctccgaaagtttccttttatggcgaggcggcggcggcggcggccctataaaaagcgaagcgcgcggcgggcgggagtcgctgcgcgctgccttcgccccgtgccccgctccgccgccgcctcgcgccgcccgccccggctctgactgaccgcgttactaaaacaggtaagtccggcctccgcgccgggttttggcgcctcccgcgggcgcccccctcctcacggcgagcgctgccacgtcagacgaagggcgcagcgagcgtcctgatccttccgcccggacgctcaggacagcggcccgctgctcataagactcggccttagaaccccagtatcagcagaaggacattttaggacgggacttgggtgactctagggcactggttttctttccagagagcggaacaggcgaggaaaagtagtcccttctcggcgattctgcggagggatctccgtggggcggtgaacgccgatgatgcctctactaaccatgttcatgttttctttttttttctacaggtcctgggtgacgaacag | PRs, RPE, inner retina, RGC | PMID: 32775493 |
| hGRK1 | atggaaaattccgagaagactgaagtggtgctgttggcatgtggatcgttcaaccccatcaccaacatgcatctgcgcctctttgaacttgccaaagactacatgaatggaactggaagatacactgtggtcaaaggcatcatctccccagtgggggatgcatacaagaagaagggcttgatccctgcctaccaccgggtcatcatggctgagctggctaccaagaactcaaaatgggtggaagtggacacctgggagtcactgcaaaaggagtggaaggaaacccttaaagtcctgcggcatcaccaggaaaagctggaagcctcggactgtgaccaccagcagaacagccccacccttgaacgcccagggagaaagcgcaagtggactgagacccaagactcaagccagaagaagtcgctggaacccaagaccaaagctgtcccaaaggtgaaactcctctgtggagctgacctcctggaatcgtttgctgtgcctaatctctggaagtcggaagatatcacccaaattgtggctaactatggcctgatctgtgtgactagagctggtaatgatgcccagaaattcatctatgaatcagatgtgctgtggaagcaccggagcaacatccatgtggtcaatgagtggattgcaaatgacatctcctccaccaagatcagaagggccctgaggcggggacagtcgatcaggtacttggtcccagaccttgtccaagagtacattgaaaagcacaacctctacagctcagagtcagaggatcgcaatgcaggagtgatcctggcccctctccagcggaacactgcagaagctaagacatag | PRs and RPE | PMID: 20428215 |
| ProC1 | agatctattagtaaaattaactacacctggtcctttatgtcattaactacacgtcagtcgttctattattaactacacagaacttactatataattaactacaccgcgcatttggatatattaactacacaactttgcttaataaattaactacacccgtaacggattctcattaactacacgtttcattacgagggattaactacacatagcccccggaggcattaactacactttttgaggtcgcgaattaactacacttcgactcgcaggacattaactacactataccgataacgcaattaactacacaaattttttatgaagattaactacactgttcgctcctgcctattaactacacgaggccgtatttccaattaactacacccgagacacaagataattaactacacttaaccagatggcagattaactacacgtacgggcaggcccgattaactacaccggctgtcattcctcattaactacacatatcacgacttgtgattaactacacgctcgagatctgcgatctgcatctcaattagtcagcaaccatagtcccgcccctaactccgcccatcccgcccctaactccgcccagttccgcccattctccgccccatcgctgactaattttttttatttatgcagaggccgaggccgcctcggcctctgagctattccagaagtagtgaggaggcttttttggaggcctaggcttttgcaaa | PRs | PMID:  31285614 |
| VMD2 | caattctgtcattttactagggtgatgaaattcccaagcaacaccatccttttcagataagggcactgaggctgagagaggagctgaaacctacccggcgtcaccacacacaggtggcaaggctgggaccagaaaccaggactgttgactgcagcccggtattcattctttccatagcccacagggctgtcaaagaccccagggcctagtcagaggctcctccttcctggagagttcctggcacagaagttgaagctcagcacagccccctaacccccaactctctctgcaaggcctcaggggtcagaacactggtggagcagatcctttagcctctggattttagggccatggtagagggggtgttgccctaaattccagccctggtctcagcccaacaccctccaagaagaaattagaggggccatggccaggctgtgctagccgttgcttctgagcagattacaagaagggaccaagacaaggactcctttgtggaggtcctggcttagggagtcaagtgacggcggctcagcactcacgtgggcagtgccagcctctaagagtgggcaggggcactggccacagagtcccagggagtcccaccagcctagtcgccagacc | RPE |  |
| ProD5 | atcctggggtttaaacaggactaggtccactgggagaagaggaagtcaaacaggactagtcatcgataggctagccagcttaattaatcagatttccaggaagaggcaccagagccctctccctattgctcaaagcgcaggtggaataatcttggccgtgtgaccggaattcaaagcagttttgggaactgagccccaacctgtaagccctgatgttatttcaagaaagatggtgtcagaattaaattggattagagaggaggatatggattagagcagagcagcatcctcacatggtcctgctggagttagtagagggta | Rods | PMID:  31285614 |
| ProC3 | Aaagctggaactggggccggaaacagctgagggtgcccagccggaaactcgaaatcaacgtaggccggaaactattcgatgaattccgccggaaactaggtccggaggactgccggaaacacctgtaatcaagccgccggaaacctgttgtggccgtatgccggaaacgtcttaattggacgtgccggaaactcttttaatgagttcgccggaaaccagaccagccgagctgccggaaaccggttatatagaacggccggaaacggtccacaggaaaaagccggaaacacccaaacggttagcgccggaaacgactggggaggacgtgccggaaacgtactatctgaagatgccggaaacacttgaaagctccaagccggaaacgggcccgtgcggatagccggaaactgacggtacacggccgccggaaacactacttgtatggtagccggaaacttgggcgtggctggggccggaaacgctcgagatctgcgatctgcatctcaattagtcagcaaccatagtcccgcccctaactccgcccatcccgcccctaactccgcccagttccgcccattctccgccccatcgctgactaattttttttatttatgcagaggccgaggccgcctcggcctctgagctattccagaagtagtgaggaggcttttttggaggcctaggcttttgcaaa | RGC,  some PRs | PMID:  31285614 |
| mOPs | Gtcaagtgagccattgtcagggcttggggactggataagtcagggggtctcctgggaagagatgggataggtgagttcaggaggagacattgtcaactggagccattgtggagaagtgaatttagggcccaagggttccagtcgcagcctgaggccaccagactgacatggggaggaattcccagaggactctggggcagacaagatgagacaccctttcctttctttacctaagggcctccacccgatgtcaccttggcccctctgcaagccaattaggccccggtggcagcagtgggattagcgttagtatgatatctcgcggatgctgaatcagcctctggcttagggagagaaggtcactttataagggtctggggggggtcagtgcctggagttgcgctgtgggagccgtcagtggctgagctcgccaagcagccttggtctctgtctacgaagagcccgtggggcagcctc | PRs | PMID: 20428215 |
| IRBPeGNAT2p | gatgtgaccatatgtcacttgagcattacacaaatcctaatgagctaaaaatatgtttgttttagctaattgacctctttggccttcataaagcagttggtaaacatcctcagataatgatttccaaagagcagattgtgggtctcagctgtgtagagaaagcccacgtccctgagaccaccttctccagctgcctactgaggcacacactgctctctcacctgccatcgtacagaccagcttttaggggagccaagttgggatactcaatcccaacttttttccttctcttccatctcacatacaggaaaccttacgagagaggattaggggcctgaaaaagctgacaagacggcaaatatgggaagtggagccagtgctgaggacaaagaactggccaagaggtccaaggagctagaaaagaagctgcaggaggatgctgataaggaagccaagactgtcaagctgctactgctgggtgagtga | Sparce cones | PMID: 24664760 |
